# Modeling stimulus-dependent variability improves decoding of population neural responses

**DOI:** 10.1101/146415

**Authors:** Abed Ghanbari, Christopher M. Lee, Heather L. Read, Ian H. Stevenson

## Abstract

Neural responses to repeated presentations of an identical stimulus often show substantial trial-to-trial variability. How the mean firing rate varies in response to different stimuli or during different movements (tuning curves) has been extensively modeled in a wide variety of neural systems. However, the variability of neural responses can also have clear tuning independent of the tuning in the mean firing rate. This suggests that the variability could contain information regarding the stimulus/movement beyond what is encoded in the mean firing rate. Here we demonstrate how taking variability into account can improve neural decoding. In a typical neural coding model spike counts are assumed to be Poisson with the mean response depending on an external variable, such as a stimulus or movement. Bayesian decoding methods then use the probabilities under these Poisson tuning models (the likelihood) to estimate the probability of each stimulus given the spikes on a given trial (the posterior). However, under the Poisson model, spike count variability is always exactly equal to the mean (Fano factor = 1). Here we use two alternative models - the Conway-Maxwell-Poisson (CMP) model and Negative Binomial (NB) model - to more flexibly characterize how neural variability depends on external stimuli. These models both contain the Poisson distribution as a special case but have an additional parameter that allows the variance to be greater than the mean (Fano factor >1) or, for the CMP model, less than the mean (Fano factor <1). We find that neural responses in primary motor (M1), visual (V1), and auditory (A1) cortices have diverse tuning in both their mean firing rates and response variability. Across cortical areas, we find that Bayesian decoders using the CMP or NB models improve stimulus/movement estimation accuracy by 4-12% compared to the Poisson model. Moreover, the uncertainty of the non-Poisson decoders more accurately reflects the magnitude of estimation errors. In addition to tuning curves that reflect average neural responses, stimulus-dependent response variability may be an important aspect of the neural code. Modeling this structure could, potentially, lead to improvements in brain machine interfaces.

## Introduction

To understand how neural responses are related to external stimuli or movements, a common approach is to characterize the average spiking activity of a neuron over repeated trials. However, neural responses show substantial trial-to-trial variability (Faisal, Selen, & Wolpert, 2008; Renart & Machens, 2014; Stein, Gossen, & Jones, 2005) – typically quantified by the Fano factor (FF): the ratio between the trial-to-trial variance and mean of the activity during a specific window of time. Although many studies assume that neural responses are Poisson distributed, with FF=1, there is strong evidence that neurons are frequently over-dispersed (FF>1) (Baddeley et al., 1997; Lehky & Sereno, 2007) or under-dispersed (FF<1) (DeWeese, Wehr, & Zador, 2003; Gur & Snodderly, 2006; Kumbhani, Nolt, & Palmer, 2007). Moreover, the variability of neural responses changes with task engagement (von Trapp, Buran, Sen, Semple, & Sanes, 2016), attention (Abolafia, Martinez-Garcia, Deco, & Sanchez-Vives, 2013), decision-making (A. K. Churchland et al., 2011), and arousal state (Lombardo, Macellaio, Liu, Palmer, & Osborne, 2018). Recent studies have also found that, even when attention and brain state do not change, variability can be stimulus-dependent (Lombardo et al., 2018; Ponce-Alvarez, Thiele, Albright, Stoner, & Deco, 2013). Altogether, trial-to-trial variability in neural responses appears to be both highly non-Poisson and also dependent on external variables. Here, using experimental data from three different brain regions, we aim to determine to what extent stimulus/behavior-dependent changes in response variability affect the ability to estimate or decode external variables.

Decoding algorithms assess to what extent the activity of a population of neurons can be used to estimate an external variable. These methods are the basis for many brain-machine interface applications (Lebedev, 2014; Schwartz, Cui, Weber, & Moran, 2006), but in most cases they may not fully capture information contained in neural variability (Quian Quiroga & Panzeri, 2009). Since neural variability is often non-Poisson and stimulus-dependent, trial-to-trial variability can carry information regarding the stimulus/behavior beyond what is accounted for under the Poisson model or any model that assumes a fixed mean-variance relationship. Models that more accurately describe trial-to-trial variability separately from the mean response could improve decoding performance. Here we examine Bayesian decoders, where we first model the likelihood of spiking activity for each individual neuron given an external variable, and we then use Bayes’ rule to compute a probability distribution over the external variable given the observed population spiking activity (the posterior). Bayesian decoding algorithms that assume neural activity is Poisson take the probabilistic nature of neural responses into account and often out-perform other methods (Chen, 2013; K Zhang, Ginzburg, Mcnaughton, & Sejnowski, 1998). However, the assumption that the response mean and response variance are equal may lead Bayesian decoders making the Poisson assumption to misestimate the posterior probability of an external variable given the observed spikes.

Here we demonstrate how the Poisson assumption can result in under- or over-confidence about external variables when decoding cortical responses, and we propose a flexible approach that allows the stimulus-dependent structure in both the mean and variance to be modeled. To account for deviations from the Poisson assumption, several previous studies have considered models with fixed, non-Poisson relationships between the mean and variance (Charles, Park, Weller, Horwitz, & Pillow, 2018; DeWeese et al., 2003; Goris, Movshon, & Simoncelli, 2014). Here we use the Conway-Maxwell-Poisson (CMP) and Negative Binomial (NB) distributions to account for non-Poisson variability and, additionally, to flexibly model how variability depends on external variables (Stevenson, 2016). These models contain the Poisson distribution as a special case, but, have an additional dispersion parameter that allows over- or, in the case of CMP, over- and under-dispersed data. Both models belong to the exponential family of distributions and can be written in the generalized linear model (GLM) framework with stimulus-dependent mean and dispersion (Sellers, Borle, & Shmueli, 2012). Here we examine three datasets: 1) recordings from primary motor cortex (M1) of a monkey performing a center-out reaching task, 2) recordings from primary visual cortex (V1) of a monkey presented with drifting sine-wave gratings, and 3) recordings from primary auditory cortex (A1) of rats presented with different pure-tone sounds. The neural responses in M1 (as a function of reach direction), V1 (as a function of grating direction) and A1 (as a function of tone frequency) have diverse tuning in both their mean firing rates and response variability. These tuning patterns can be accurately described by CMP and NB models; moreover, in these three cortical areas, we find that Bayesian decoders using the CMP or NB models improve estimation by 4-12% compared to the Poisson model. The additional layer of information in stimulus-dependent variability thus appears to improve decoding performance.

## Methods

### Neural Data

Here we use 3 datasets recorded from primary motor (M1), visual (V1), and auditory (A1) cortices to determine whether modeling dispersion improves decoding accuracy.

For the primary motor cortex, we used the DREAM-Stevenson_2011 dataset from CRCNS (Walker & Kording, 2013). This dataset was recorded from the arm area of primary motor cortex of an adult macaque monkey during center-out reaches. The reaches were made in a 20×20 cm workspace while the animal was grasping a two-link manipulandum. The data was recorded using a 100-electrode Utah array (400 mm spacing, 1.5 mm length) and was spike sorted manually with an offline sorter (Plexon, Inc) where 69 single units were identified with >.2 spikes per trial on average. On each trial, we analyzed spike counts during the window 150 ms before to 350 ms after the speed reached its half-max. Detailed descriptions of the surgical procedure, behavioral task, and preprocessing are available in the original report (Stevenson, 2016).

For the primary visual cortex, we used the PVC-11 dataset from CRCNS (Kohn & Smith, 2016). This dataset was recorded from an anesthetized adult monkey (monkey 3) in response to presentations of drifting sine-wave gratings (with 20 trials for each of 12 directions). The data was recorded using a 96-channel multielectrode array and 112 single units were identified with SNR >1.5 and firing rate >1 Hz. On each trial we analyzed spike counts between 200 and 1200 ms after stimulus onset. Detailed descriptions of the surgical procedure, stimulus presentation, and preprocessing are available in the original reports (Kelly, Smith, Kass, & Lee, 2010; Smith & Kohn, 2008).

For primary auditory cortex, in contrast with M1 and V1 data that were simultaneously recorded, we combined extracellular recordings from 18 anesthetized male rats. These data were recorded from primary (A1; n=176) auditory area using 16 channel tetrodes. Spikes were detected and clustered using custom routines and later sorted with automated clustering (KlustaKwik). Neurons were responding to sound stimuli for measuring frequency response areas (FRA) with transient tones over a range of frequency (1.4-45.3 kHz; 42 frequencies with one-eighth-octave steps) and sound pressure levels (85 dB to 15 dB in 10-dB steps). On each trial, we analyzed spike counts during a 100 ms window after the stimulus onset. Each combination of frequency and sound level was presented six times in random order. Here we only use single units from area A1 with >.1 spikes per trial on average (n=158). We decode stimulus frequency after re-binning the stimuli into 21 categories, and we combine responses from three sound pressure levels (45, 55, and 65 dB) to get 36 “repetitions” per category. Note that, since trials at the same frequency contain different sound levels, there may be higher than expected variability in neural responses. Detailed descriptions of the surgical procedure, stimulus presentations, and preprocessing are available in the original report (Lee, Osman, Volgushev, Escabí, & Read, 2016).

### Bayesian Decoding

The goal of population decoding is to take the spike counts from all neurons on each trial and to estimate what stimulus/movement occurred during that trial. Bayesian decoding methods make use of the fact that the relationship between an external variable θ and population neural activity **N**_θ_ = [**n**_1_(θ)…**n**_C_(θ)] can be modeled with Bayes rule by *p*(θ|**N**_*θ*_) ∝ *p*(**N**_*θ*_|θ)*p*(θ) (Chen, 2013). Where the posterior distribution *p*(θ|**N**_*θ*_) over the external variable is determined by the product of the likelihood *p*(θ|**N**_*θ*_), the probability of observing the given neural activity under a specific encoding model, and the prior *p*(θ) that determines how likely each value of the external variable is, *a priori*. In most applications of Bayesian decoding, the neurons are assumed to be conditionally independent given the external variable: *p*(**N**_*θ*_|θ) = Π_*i*_*p*(**n**_i_(θ)|θ). This assumption allows straightforward estimation of the posterior, but, in practice, modeling dependencies between neurons can improve decoding (Pachitariu, Petreska, & Sahani, 2013; Park, Archer, Latimer, & Pillow, 2013; Stevenson et al., 2012). Here we use the independence assumption and examine how the choice of likelihood affects decoding performance.

Previous works have almost exclusively considered Poisson likelihood models:

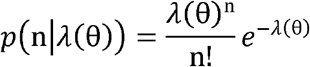

where n denotes the observed spike count and *λ* is a mean spike count that changes as a function of the external variable, θ. Here we use cubic basis splines, b_k_(θ), with equally-spaced knots to describe the tuning curves of each individual neuron: 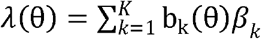 with K=5, 6, and 8 for M1, V1, and A1, respectively. When θ is a circular variable, the splines are assumed to have periodic boundary conditions. In order to estimate the coefficients, ***β***, we assume observations are conditionally independent and impose an L1-penalty on the coefficients to reduce overfitting:

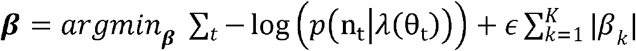

where *t* indexes the individual external variable on each trial. Once we have estimated the tuning curve *λ_i_*(θ) for individual neurons in our training set, we apply Bayes’ rule to decode the stimuli/movement for each trial in the test set (8-fold cross-validation). The hyper-parameter, *ϵ*, was selected with a line search for each neuron. In general, we assume that the prior *p*(θ) in our decoders is flat and each stimulus is equally probable.

### Conway-Maxwell-Poisson Models

In previous work we described how CMP models can provide more accurate descriptions of trial-to-trial variability for tuning curves (Stevenson, 2016). The CMP distribution takes the form:

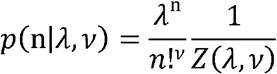

with normalization factor 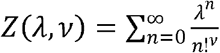. For spike counts n, the distribution is a function of the intensity, *λ*, and dispersion parameters, *υ*, with *υ* > 1 describing over-dispersion and *υ* > 1 describing under-dispersed data. Since there is no known closed-form solution for the normalization factor (Minka, Shmueli, Kadane, Borle, & Boatwright, 2003), we instead compute it numerically up to some finite sum. Note that with *υ* = 1, *Z*(*λ, υ*) = *e^λ^* and the CMP is exactly the Poisson distribution, and the spike counts are equi-dispersed.

In practice, we take advantage of the fact that the CMP distribution is in the exponential family and frame the problem of tuning curve estimation as a generalized linear model (GLM) (Sellers et al., 2012). In particular, we estimate parameters ***β*** and ***γ*** that map external covariates **ϔ**(θ) = [b_1_(θ) … b_k_(θ)] and **Ψ**(θ) = [c_1_(θ) … c_L_(θ)] to neural responses using the link functions log(*λ*(θ)) = ϕ(θ)***β*** and log(*υ*(θ)) = **Ψ**(θ)***σ***. This framework is in effect a dual-link GLM where both the mean and the variance depend on the external variable θ (Sellers & Shmueli, 2010).

We again estimate the tuning curves using spline bases and maximum a posteriori (MAP) estimation, here with L2 regularization. Importantly, this approach allows us to model neural responses that are under-dispersed, over-dispersed, or that contain both under- and over-dispersed counts in response to different stimuli or movements. As with the Poisson models, once we have the likelihood of spike responses, we use a Bayesian decoder to estimate the posterior distribution over external variable given spiking: *p*(θ|**N_θ_**) ∝ *p*(**N_θ_**|θ)*p*(θ) and assume that the neurons are conditionally independent.

We also compare the Poisson with the Negative Binomial model which has been used more widely to describe over-dispersed spike counts (Scott & Pillow, 2012; Taouali, Benvenuti, Wallisch, Chavane, & Perrinet, 2016):

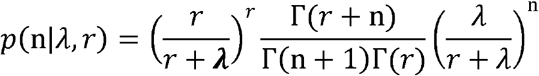

where the NB distribution is parameterized by the mean *λ* and dispersion parameter *r*, and Γ(·) denotes the gamma function. The NB model can also be written as canonical generalized linear models using the link function log(***λ***(θ)) = **ϕ**(θ)***β***, and, as with the CMP model, covariate-dependent dispersion can be modeled by assuming log(*r*(θ)) = **Ψ**(θ)***γ***. With both the CMP and NB models we use L=3, 4, and 6 for the M1, V1, and A1 datasets, respectively. Altogether, these three models (Poisson, NB, and CMP) allow us to describe a range of tuning behaviors in different brain regions and to quantify to what extent modeling stimulus/movement-dependent dispersion improves decoding accuracy.

### Linear Decoding

In addition to comparing our non-Poisson Bayesian decoders to the conventional Poisson Bayesian decoder, we also compare these models to two non-probabilistic linear decoding methods. Linear decoders estimate the external variable *θ* on each trial using a linear combination of neural responses and weight functions ***Ψ***(*θ*). This assumption is the basis for population vector approaches (Georgopoulos, Schwartz, & Kettner, 1986), as well as, template matching (Wilson & McNaughton, 1993), and optimal linear estimation (OLE) methods (Salinas & Abbott, 1994). Here we examine template matching and OLE. For template matching we estimate the tuning curves for each neuron ***Ψ***_*i*_(*θ*), by linear (ridge) regression using K cubic basis spline functions, **ϕ**(θ) ***Ψ***_*i*_(*θ*) = **Σ**_*k*_ ***b***_*i,k*_ **ϔ**_k_ : θ). Namely, we estimate weights ***b*** = *b_i,k_* = by

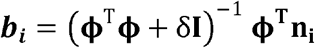

where **n_i_** = n_i,t=1:T_ is a vector constructed from the neural response of neuron *i* on trial *t*. **ϕ** = **ϕ**_k=1:K,t=1:T_ is a matrix representation of the external variable formed by basis functions. As with the tuning curves for Bayesian decoding described above, we use cubic B-Spline bases with equally-spaced knots and periodic boundary conditions where appropriate (V1 and M1). δ is a regularization hyperparameter to prevent overfitting which we select by grid search over logarithmically-spaced values from .001 to 10 and minimizing the cross-validated squared error. Given the tuning curves for each neuron, we then evaluate the performance of our template matching decoder by constructing linear estimates of the external variable 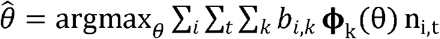.

For optimal linear estimation (OLE), rather than fitting tuning curves for each neuron independently, we estimate weights, *b*, for all neurons, simultaneously. Using a similar notation as above,

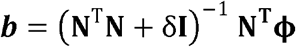

where **N** is now the spike count matrix for all neurons. As with template matching, estimates of the external variable are linear. However, the weight matrix here differs from the one in template matching as OLE accounts for correlations in the population responses.

### Parametric accuracy and error curves

When estimating the decoding accuracy or error as a function of the number of neurons included in the model, we often cannot exhaustively evaluate all possible 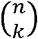 subpopulations. For clarity, we thus fit a parametric curve to accuracy and error estimates as a function of population size k. We define a curve that follows a generalized logistic function:

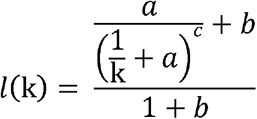

with the parameters {*a*, *b*, *c*}. We then fit *l*(k) to accuracy curves and *h*(k) = 1 − *l*(k) to error curves for random subsamples of the full set of neurons by minimizing the squared-error and constraining the parameters to be positive.

### Greedy algorithm

In addition to quantifying the decoding performance of a random set of neurons, it is also useful to consider the best and worst performing sets of k neurons. These extremes give some indication as to what the limits of neural coding might be. To find these subsets of neurons that give upper or lower bounds of the accuracy, we use a greedy combinatorial optimization scheme similar to beam search. Namely, to maximize decoding accuracy, we start with an empty set, then we add neurons one-by-one and select the best set at each stage. To avoid local minima, we keep the top-5 performing sets at each stage and consider additions to each of them, rather than only keeping the top performing set (usually called a “greedy” search). To find the worst performing set of neurons, we simply keep the worst sets rather than the best. This approach is not guaranteed to strictly maximize/minimize the decoding accuracy over all possible 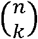 subsets, but it provides approximate limits on performance.

For comparison we also consider taking the neurons with the best/worst tuning. We identify the best tuned neurons by selecting those with the lowest tuning index: 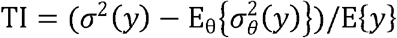 where E{y} and *σ*^2^(y) are the total mean and variance of the responses and 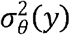 is the variance for trials with stimulus/movement θ.

## Results

Traditionally, spike count models assume Poisson distributed responses where the average neural response across multiple trials of a specific external variable equals the variance in the neural responses across trials. However, observations from different areas and brain states show that neural responses often deviate from the Poisson assumption (Baddeley et al., 1997; Gur & Snodderly, 2006; Kumbhani et al., 2007; Lehky & Sereno, 2007; Lombardo et al., 2018; Ponce-Alvarez et al., 2013). Here we demonstrate how flexible models of stimulus- and movement-dependent dispersion can improve neural population decoding.

To illustrate how neural variability can alter estimates of external variables we consider Bayesian decoding (Chen, 2013; Kechen Zhang & Sejnowski, 1999) where we aim to evaluate the posterior probability of the external variable given observations of neural activity. The central insight for Bayesian decoding is that this posterior can be written (following Bayes’ rule) as the product of a likelihood (the probability of neural responses given an external variable) and a prior (probability distribution over the external variable, see Methods). Although most previous work with Bayesian decoders has assumed that the likelihood follows a Poisson distribution, assuming Poisson noise when in fact spike counts are under- or over-dispersed can thus result in under- or over-confidence in the decoding, respectively. Generally, stimulus-dependent changes in the variability will affect the shape of the posterior distributions (Fig. 1). If the dispersion varies as a function of the stimulus dimension distinctly from the mean, the maximum of the posterior can shift and change the stimulus estimate (Fig 1A, right). Additionally, neurons with flat tuning curves, that would generally be considered uninformative, can provide information about the stimulus if they have stimulus-dependent variability (Fig. 1B).

**Fig. 1:**
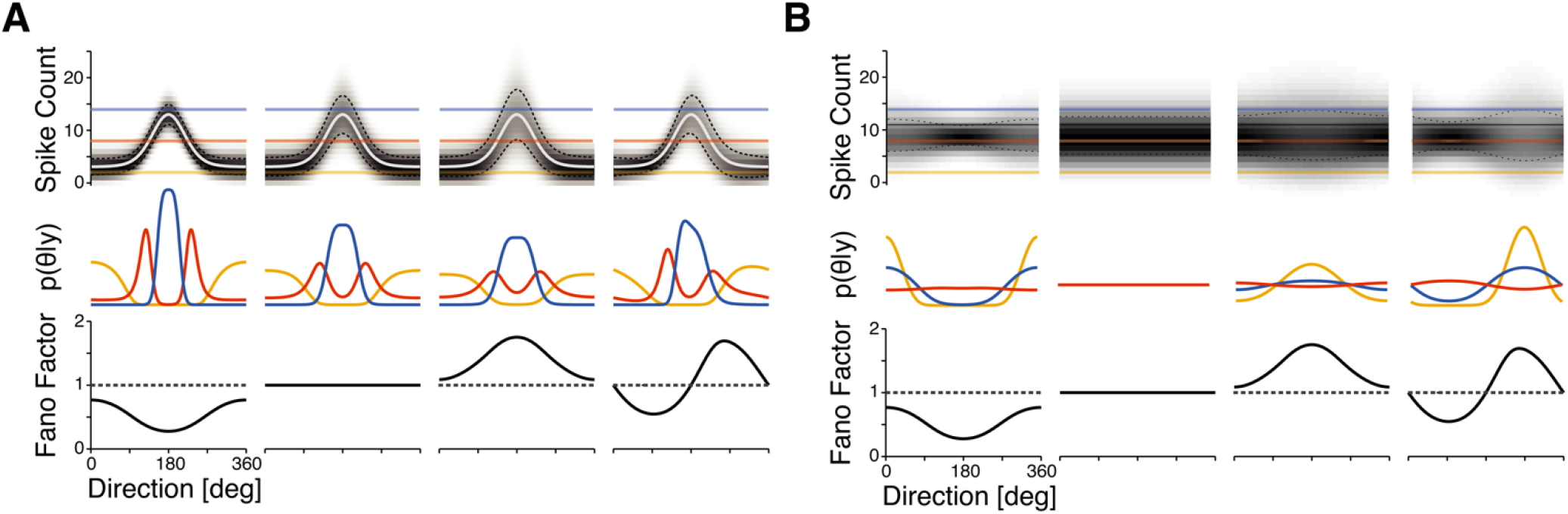
Stimulus-dependent variability alters posterior probabilities for sensory or motor variables. A) Four simulated neurons with identical mean tuning curves and different types of dispersion. From left to right, spike counts are under-dispersed, equi-dispersed, over-dispersed, and both under- and over-dispersed depending on the stimulus. Note that although the encoding distributions (top) are similar, the decoding distributions (middle) show substantial differences. Blue, red, and yellow curves indicate the posteriors over stimuli/movement direction with observations of 14, 8, and 2 spikes on a trial, respectively. Increased spike count variability leads to increased uncertainty in the stimulus, and if the mean-variance relationship is not fixed then the maxima of the posterior may also be different (right-most neuron). B) Neurons that would typically be called “untuned”, based on their mean responses, can provide stimulus/movement information if the dispersion is stimulus/movement-dependent. Fano factors profiles for all four neurons are the same as A.

In experimental data from primary motor (M1), visual (V1), and auditory (A1) cortex (see Methods), we find that in addition to traditional tuning of the mean, many neurons also have tuning in dispersion (Fano factors). The correlation between mean firing rate and Fano factor tuning curves spans the whole range from −1 to 1 (Fig. 2) with average correlations in M1: 0± 0.5, V1: 0.2±0.4, and A1: 0±0.3 SD. The Fano factors themselves also vary over a wide range in each of these areas (M1: 1.6± 0.6, V1: 2.8±1.3, and A1: 1.5±0.5, mean±SD). Rather than a uniform population of neurons with approximate Poisson firing (Fano factor near 1), we find a wide diversity of stimulus and movement-dependent patterns in the Fano factors. Although these diverse patterns of stimulus-dependent variability are not well described by the Poisson model, they are well described by the non-Poisson, CMP model with variable dispersion (fits in Fig. 2B, see Methods).

**Fig. 2:**
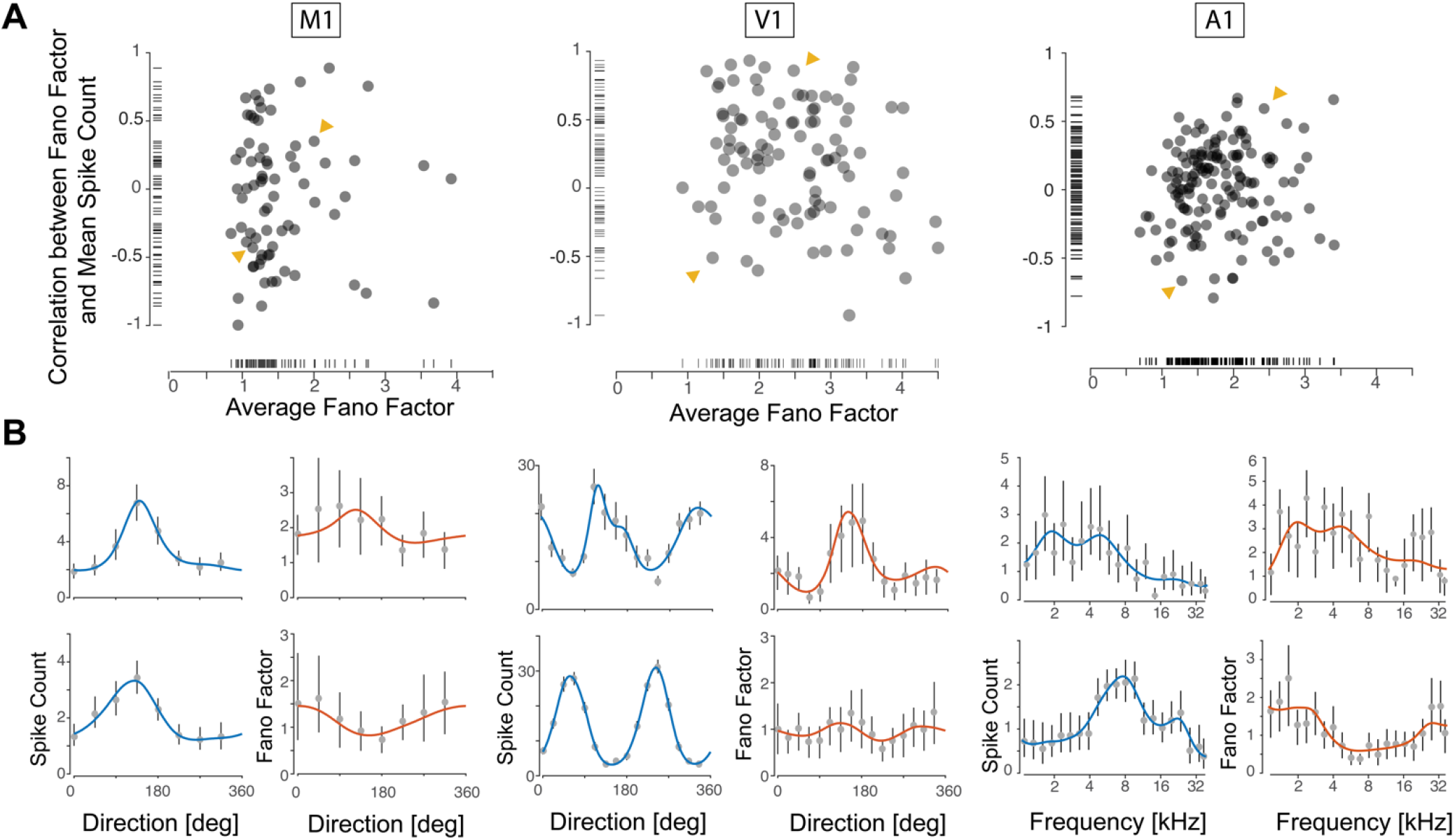
Diversity in tuning curve dispersion. A) Fano factors and correlation between Fano factor and mean spike count tuning curve for M1, V1, and A1. Note that the Fano factors are not well described as constant close to 1 (as would be the case for Poisson firing) or even well correlated with the spike count (as would be the case for a fixed, increasing mean-variance relationship). B) Two example neurons from each area are shown, corresponding to the arrows in A. Curves show fits from CMP models with stimulus/movement-dependent dispersion; dots and error bars denote observed means and 95% confidence intervals (estimated by Bayesian bootstrapping, see (Rubin, 2007)).

For each neuron in the three cortical areas we aligned the firing rate tuning curves from each neuron by their preferred stimulus/movement direction (Fig. 3, top). As expected, neurons in all areas show increased activity around the preferred stimulus/movement. Note that the tuning curves often contain multiple maxima, particularly in V1. Similarly, we align Fano factor profiles based on the preferred stimulus/movement in firing rate tuning curves (Fig. 3, bottom). For many neurons, the Fano profiles do not show the same alignment as the firing rate responses. This observation suggests that although both the mean and Fano factor tuning curves are stimulus-dependent, their dependencies are not necessarily matched to each other. The correlation (circular for M1 and V1) between the preferred stimulus/movement and the stimulus/movement with the maximum Fano factor tends to be low (M1:0.24±0.05, V1 :-0.03±0.02, and A1:0.11±0.01 mean±SE, bootstrapping across neurons). These low correlations seem to suggest that stimulus/movement-dependent variability is flexible, and not simply the result of a fixed, monotonically increasing mean-variance relationship.

**Fig. 3:**
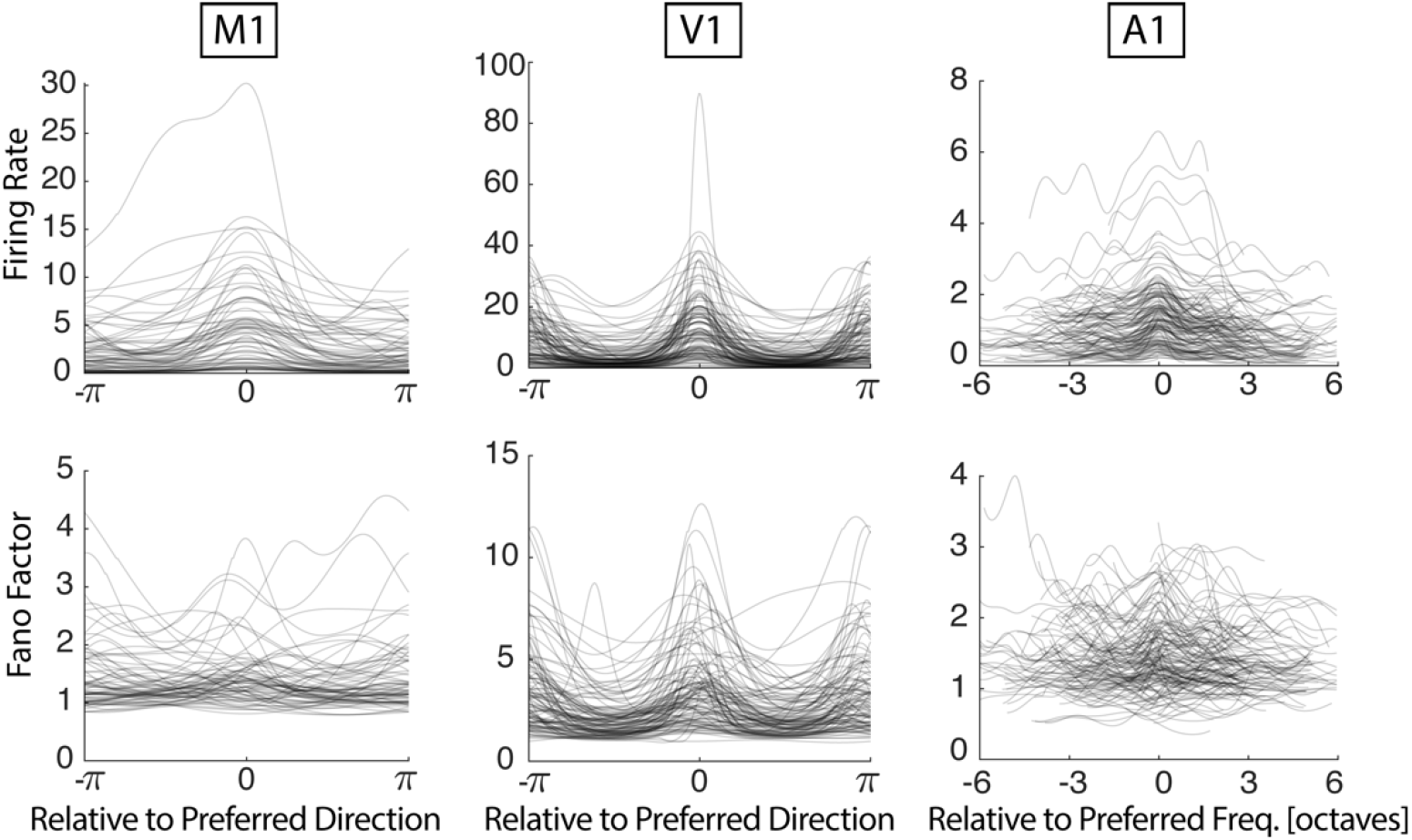
Average response and Fano factor are not necessarily aligned to the same preferred stimuli/movement. Top, response tuning curves of individual neurons aligned with their maximum firing rate. Bottom, Fano factor tunings of individual neurons aligned with the preferred stimulus from firing rate tuning curves.

To explore how these patterns of dispersion affect decoding, we compare the decoding accuracy and prediction error under different models in these three different cortical areas (Fig. 4). For all models and recording areas, decoding accuracy increases and prediction error decreases as more neurons are included in the Bayesian decoders (with Poisson, Negative Binomial, and Conway-Maxwell-Poisson likelihoods), as well as, linear decoders (using template matching and Optimal Linear Estimation). The Bayesian decoders with Negative Binomial and Conway-Maxwell-Poisson noise models generally outperform the Bayesian decoders that assume Poisson noise. These models both have the flexibility to describe stimulus/movement-dependent variability in addition to mean tuning. Taking the best performing model between the CMP and NB models, the non-Poisson models are 3.5±0.5% (M1), 3.8±0.3% (V1), and 11.8±1.7% (A1) better than the Poisson models when using the full population of neurons (mean±SE bootstrapping across trials). These improvements also imply that the same performance level can be achieved with fewer neurons. For example, to reach an accuracy of 75% in the V1 dataset the Poisson model uses 34 randomly selected neurons while the CMP only uses 27 neurons. For reference, we also evaluated the performance of two non-probabilistic, linear decoding methods: template matching and optimal linear estimation. Template matching tends to perform worse than the other methods, but OLE achieves relatively high accuracy and low prediction error. However, OLE typically does not reach the performance of the Bayesian decoders until there are many neurons in the model. Note that, although the neurons in M1 and V1 datasets were simultaneously recorded, the A1 contains neurons combined from different animals and different sessions.

**Fig. 4:**
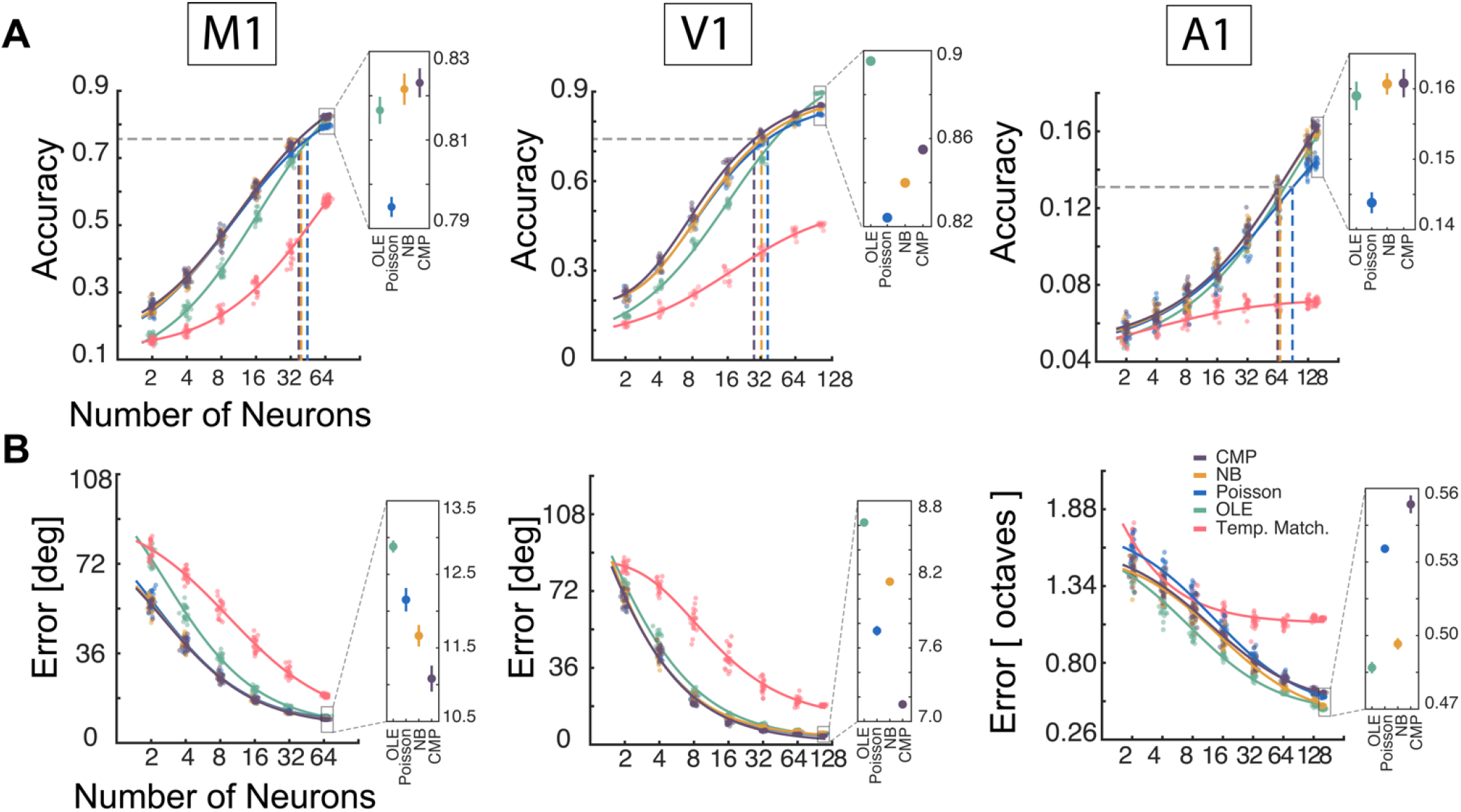
Modeling dispersion improves the accuracy of population decoding. Decoding accuracy (A) and error (B) for different population sizes in primary motor (M1), visual (V1), and auditory (A1) areas for five models: template matching (red), optimal linear estimation (green), Poisson (blue), negative binomial (yellow), and Conway-Maxwell-Poisson (purple) Bayesian decoders. Jittered data points denote the performance of random subsets of neurons, and curves denote parametric fits. The insets show the performance for the largest subset size (N-5), and error bars denote standard deviation across subsets.

Although we use the mode of the posterior in Bayesian decoders as our point estimate of the stimulus/movement, the posterior also reflects the uncertainty about the identity of the stimulus/movement under the different models. When aligned with the true stimulus/direction the three different Bayesian decoders have distinct posterior distributions (Fig. 5A). To quantify the uncertainty in the posteriors, we look at the standard deviation (SD - circular SD for direction stimuli in M1 and V1). The (circular) SD of the posterior increases with increasing absolute prediction error suggesting that trials with bigger errors have higher uncertainty (Fig. 5B). Additionally, we observe that the two non-Poisson models (NB and CMP) have wider posteriors compared to the Poisson models, for all trials. We also examined how accurately the posterior covers the true stimulus/movement. Namely, we examine how the coverage (the fraction of trials on which the true stimulus falls within a confidence region) varies as we change the confidence-level (the area of the confidence region when selecting the highest probabilities first) (Fig. 5C). For instance, if we set the confidence-level to 0.95, we would expect the true stimulus/movement to fall within confidence region 95% of the time if the model is accurate. Here we find that all models tend to be over-confident, with the Poisson model being the most over-confident. For example, in the A1 dataset a 95% confidence level covers the true stimulus for only 46% of trials in the Poisson model, while covering 59% and 62% of the stimuli in the CMP and NB models.

**Fig. 5:**
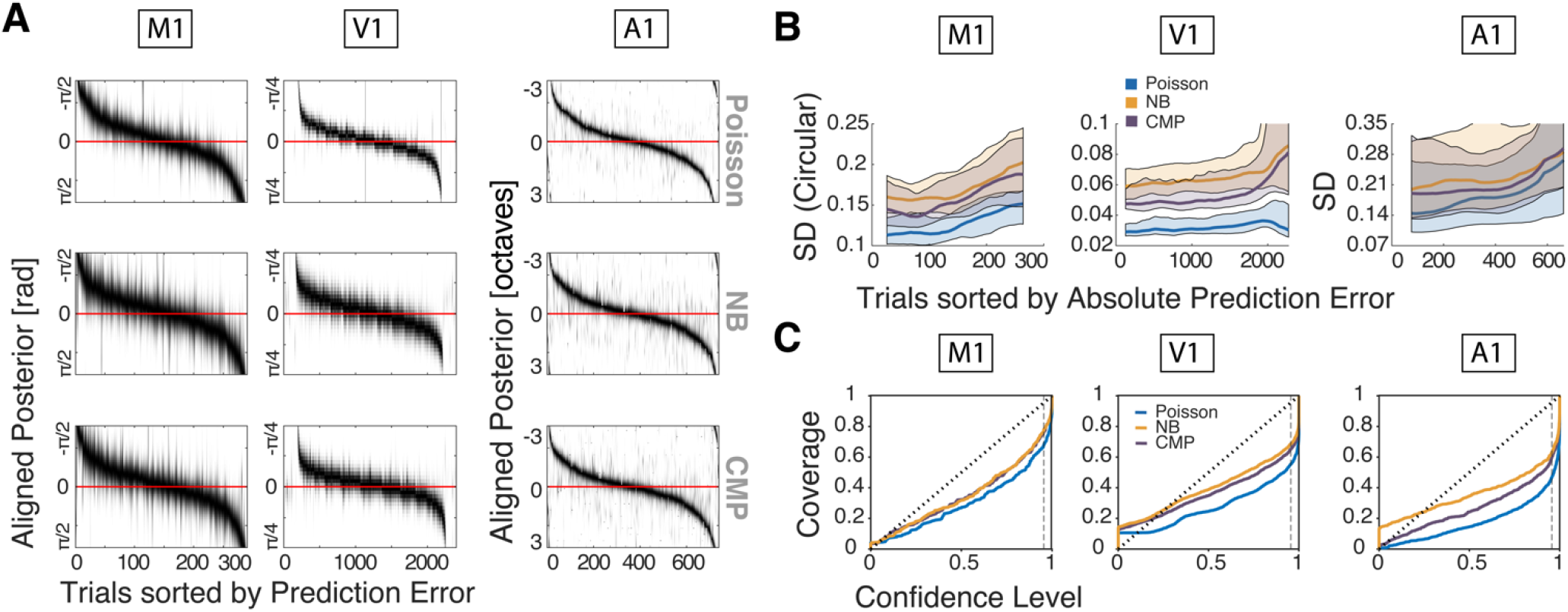
Bayesian models assuming Poisson variability tend to be over-confident. A) Decoded posterior distributions for all trials sorted by prediction error. In three primary areas (M1, V1, and A1) we aligned posterior with true stimulus (red line). Black shades show higher probability in posterior. For Negative Binomial and CMP, the posterior covers the true stimulus in more trials. B) Standard deviation of the posterior increases for all models as (absolute) prediction error increases. Additionally, the non-Poisson models tend to have higher uncertainty (SD) compared to the Poisson model, for all trials. Solid lines denote a moving median over 50 (M1), 250 (V1), and 150 (A1) trials and error bands denote inter-quartile range. C) Comparing the coverage of the three different models: the fraction of trials where the true stimulus is contained by a confidence region of a given size, i.e. the confidence level, we find that all three models are over-confident. However, the posteriors of the non-Poisson models cover the true stimulus more accurately.

Decoding performance improves when more neurons are included in the model. However, in addition to characterizing the performance of randomly selected populations of neurons, we can also examine which neurons contribute most or least to decoding performance and optimize performance by using only a select subset of neurons. To identify approximate upper-bounds for the decoding performance of a population of a given size we search for subsets of best performing neurons using a greedy algorithm: beam search (see Methods). For all three datasets, beam search identifies subsets of neurons that increase decoding accuracy well above that of random populations neurons of the same size and also populations of the same size with the best tuning. For all three datasets a greedy search can reach to certain level of accuracy with much lower number of neurons. Moreover, in all datasets, dispersion models perform similar or better than Poisson in reaching high accuracies with fewer neurons.

**Fig. 6:**
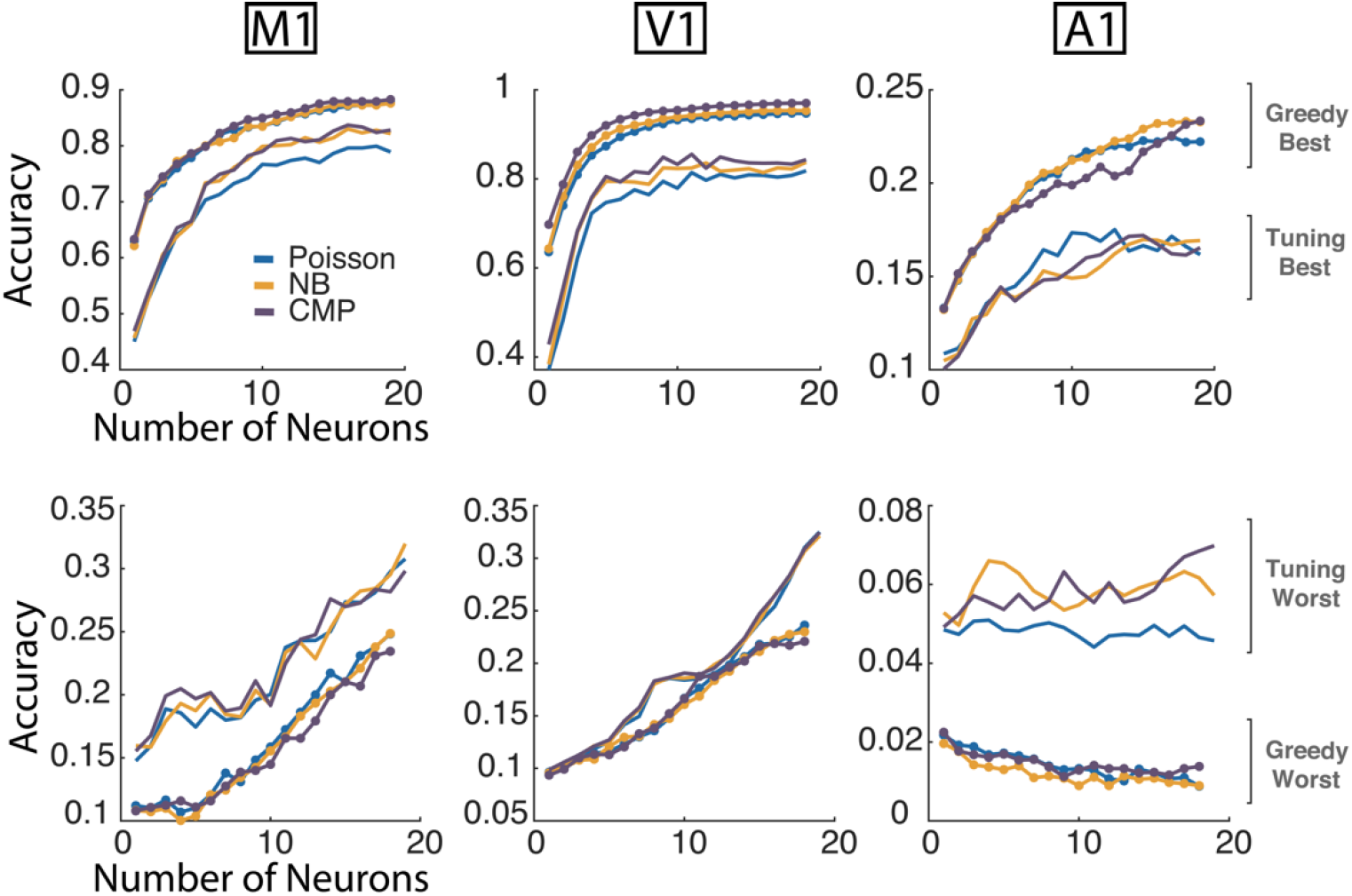
Best- and worst-case decoding performance. Decoding accuracy of the best/worst performing subpopulations of neurons (found with beam search) and the most/least tuned subpopulations. For Poisson (blue), NB (yellow), and CMP (purple) Bayes decoders the best/worst case performance tends to be similar and well above/below the performance of random subpopulations of the same size (see Fig 4). Greedy best and worst results are the average of the top-5 sets that were found.

## Discussion

Neural responses to repetitions of an identical external variable can have substantial trial-to-trial variability. Depending on the brain area and experimental setting, spike counts can show both over- or under-dispersion – variances bigger or smaller than mean, respectively. Here we model stimulus/movement-dependent changes in spike count variability and characterize to what extent this structure can be used for more accurate neural decoding. Our results show that probabilistic models that account for stimulus-dependent under- or over-dispersion in the data (Conway-Maxwell-Poisson and Negative Binomial) can improve decoding performance in M1, V1, and A1 compared to traditional Poisson-based Bayesian decoders. These results are consistent with recent work showing that Negative Binomial models can improve decoding in area MT (Taouali et al., 2016). Additionally, we find that decoders with flexible models of dispersion more accurately reflect decoding uncertainty compared to the Poisson models.

Many studies have noted that neural variability is non-Poisson and have proposed other mean-variance relationships (Amarasingham, 2006; Charles et al., 2018; Goris et al., 2014). However, in addition to being non-Poisson, results also suggest that the relationship between the mean and variance of neural responses is state and stimulus-dependent (Lee et al., 2016; Lombardo et al., 2018; Ponce-Alvarez et al., 2013) and may not be fixed. Here we aimed to determine how stimulus/movement-dependent mean and variance relationships change Bayesian decoding. In particular, to model these relationships, we use the Conway-Maxwell-Poisson and Negative Binomial models that can both be framed as generalized linear models (GLMs). By making their dispersion parameters stimulus/movement-dependent, these GLMs are flexible enough to model the wide range of mean-variance relationships observed in experimental data. In the three datasets examined here, the majority of neurons are over-dispersed and the CMP and NB models perform similarly. However, the CMP may be preferable when the data is under-dispersed, since the NB only allows for over-dispersion. At the same time, fitting the CMP model is less tractable, since evaluating the CMP log-likelihood requires calculating a computationally intensive normalization. Recent work has also introduced even more flexible count models (Gao, Buesing, Shenoy, & Cunningham, 2015), that could further improve performance. Additionally, it is important to note that here we used rate-based decoders and model spike counts on relatively long timescales, with bin sizes >100 ms in all datasets. The Fano factor depends heavily on bin size (Baddeley et al., 1997; Warzecha & Egelhaaf, 1999), and modelling the more detailed temporal structure of neural responses can improve decoding in many neural systems (Butts et al., 2007; Lee et al., 2016; Osman, Lee, Escabí, & Read, 2018).

Although modeling stimulus-dependent, non-Poisson variability can improve decoding accuracy in practice, these results may also have implications for theories of population coding. Assuming Poisson variability allowed previous theoretical studies to make strong predictions about how the shape and spacing of the tuning curves in a population of neurons could be optimized to collectively represent sensory or motor variables (Alexandre Pouget, Deneve, Ducom, & Latham, 1999; Kechen Zhang & Sejnowski, 1999). In some cases, decoding with the Poisson assumption can be equivalent to decoding using linear combinations of neural activity (Ma, Beck, Latham, & Pouget, 2006), which could hypothetically provide a simple computational basis for Bayesian behavior (A Pouget, Dayan, & Zemel, 2003). Previous studies have characterized the overall impact of variability on sensory coding (Butts & Goldman, 2006), but the diverse patterns of neural variability that we observe in data seem to suggest that probabilistic representations of stimuli and movements could be affected by structured patterns in trial-to-trial variability more broadly.

Altogether, our findings suggest that task and stimulus-dependent variability may be an important aspect of the neural code. Non-Poisson counts models such as the CMP and NB here could be used to explore the count variability and co-variability that have been linked to stimulus-onset (M. M. Churchland et al., 2010) or to attention and learning (Mitchell, Sundberg, & Reynolds, 2009). Since the CMP and NB models can be formulated as GLMs, other covariates, such as local field potentials (Niknam, Akbarian, Noudoost, & Nategh, 2017), plastic neural interactions (Ghanbari et al., 2018), or even latent variables (Chase, Schwartz, & Kass, 2010; Kulkarni & Paninski, 2007; Lawhern, Wu, Hatsopoulos, & Paninski, 2010), can easily be included in the models. Ultimately, the CMP and NB models provide a framework to describe the stimulus-dependence of both the mean and variance of neural responses. Here we show how modeling this dependence can improve decoding.

